# Older adults do not have a higher metabolic cost than younger adults in outdoor overground walking

**DOI:** 10.64898/2026.07.27.740899

**Authors:** Eline van der Kruk, Koen Jongbloed, Monica Orlandi, Michael F. Miller, Anne K. Silverman

## Abstract

The metabolic cost of walking is widely used to evaluate human performance and effectiveness of clinical interventions. Decades of laboratory research, largely based on treadmill experiments, have established a canonical relationship between walking speed and metabolic cost, and suggested that ageing shifts this relationship upward, implying reduced efficiency in older adults. However, this relationship has not been well tested during overground walking across matched speeds. We compared healthy younger (n=16; 26±2yr) and older (n=11; 74±3yr) adults across eight outdoor overground walking trials at different speeds: preferred walking speed (PWS), three fixed speeds (0.8, 1.2, 1.6 m·s⁻¹), and four speeds at ±5% and ±10% of PWS. Contrary to our hypothesis, older adults did not show higher gross or net metabolic cost of walking (GCOW and NCOW) than younger adults at any speed; rather, both trended consistently lower in older adults, reaching significance for GCOW at 0.8 m·s⁻¹ only. Comparisons of resting metabolic rate and respiratory exchange ratio to prior reference groups did not indicate that our older cohort was unusually fit. Independent of age, GCOW was significantly higher at 0.8 m·s⁻¹ than at the remaining speeds (1.2–1.6 m·s⁻¹), confirming that walking at slower speeds increases GCOW. These findings challenge the view that ageing intrinsically increases the energetic cost of walking, suggesting instead that previously reported upward shifts in cost may reflect treadmill-specific constraints or speed effects. Future work is needed to explore direct comparisons of outdoor, overground walking with treadmill walking at fixed speeds in both younger and older adults.

## Introduction

Metabolic cost of walking (MCOW), defined as metabolic energy expended per meter travelled, is often used to quantify human performance (Boyer et al., 2023), to compare the efficacy of interventions (e.g., Galle et al., 2017; Montgomery & Grabowski, 2018), and to develop and validate musculoskeletal movement simulations (Koelewijn et al., 2018; Miller et al., 2024; Umberger, 2010; Veerkamp et al., 2021). The relationship between MCOW and walking speed has been widely studied, with the minimum cost of walking often observed at a self-selected walking speed, also known as preferred walking speed (PWS) when no specific instructions about speed are given. Many studies consider this relationship to be parabolic (Larish et al., 1988; P. E. Martin et al., 1992) or hyperbolic (Ralston, 1958), with the hyperbolic relationship resulting from a group of individuals who have a parabolic relationship between walking speed and metabolic cost (Ralston, 1958). The MCOW-walking speed relationship, most often explored on treadmills, has become a foundational concept in gait biomechanics (Malatesta et al., 2003; P. E. Martin et al., 1992; Ralston, 1958; Zarrugh et al., 1974).

How this MCOW-walking speed relationship may change across the lifespan has long been of interest (e.g., Larish et al., 1988; Pearce et al., 1983). Older adults in biomechanics studies are regularly observed to have a lower PWS and a higher MCOW measured in laboratory environments compared to younger adults, but the mechanism behind these changes is not clear (Boyer et al., 2023; Das Gupta et al., 2019; Schrack et al., 2012). Longitudinal studies have proposed that this higher MCOW may precede a reduction in PWS (Schrack et al., 2016), which can be a marker for long term mobility and health. Assuming an underlying parabolic relationship where MCOW is lowest at a single preferred PWS, lower PWS and higher cost of walking in older adults may be explained by an upward shift of this curve or PWS selection based on criteria other than energy optimization. Older adults have altered musculoskeletal capacity and thus may weigh multiple movement objectives differently than younger adults (van der Kruk et al., 2021).

Multiple prior studies have found an upward shift in MCOW for older adults relative to younger adults, but these studies have all been on a treadmill (Larish et al., 1988; Malatesta et al., 2003; P. E. Martin et al., 1992; Zarrugh et al., 1974). However, there are several differences between overground and treadmill walking. Overground walking differs from treadmill walking in metabolic cost, spatiotemporal gait characteristics and preferred walking speed (Berryman et al., 2012; Larish et al., 1988; J. P. Martin & Li, 2017; Schmitt et al., 2021). In addition, how individuals adapt to walking surface and context is likely age dependent. That is, when walking on a treadmill, older adults have a greater increase in cost compared to younger adults at PWS compared to overground walking (Das Gupta et al., 2021), and acclimatization to treadmill walking may require substantial time (Berryman et al., 2012; Das Gupta et al., 2019). The context in which MCOW is measured is likely to affect interpretation of age-related differences.

Overground studies comparing older and younger adults are sparse. A recent meta-analysis found higher gross and net COW in older adults at PWS (Das Gupta et al., 2019), but only 2 of 19 included studies were overground (Waters et al., 1983, 1988). One of these studies only measured at PWS (Waters et al. 1983), which was slower in older adults, leaving unclear whether observed differences reflect physiological aging or speed effects. A landmark study comparing age groups walking at multiple speeds, slow, normal and fast, (Waters et al., 1988) found no difference in gross cost of walking (GCOW) between healthy adults (20-59 years) and seniors (60-80 years). However, as walking speeds weren’t matched, there was still the confounding factor of speed, making potential age-related effects difficult to isolate. Overground studies at a single matched walking speed reported no difference in MCOW between age groups (Das Gupta et al., 2021). Whether the upward shift of the MCOW-walking speed curve seen on treadmills in older adults also occurs during overground walking remains unknown.

To evaluate how MCOW is related to walking speed, a gap in our knowledge exists in an overground experiment of older adults compared to younger adults, which includes multiple matched walking speeds. To address this gap, we measured metabolic cost across multiple outdoor, overground walking speeds in younger (YA) and older adults (OA), including PWS, three fixed speeds, and ±5% and ±10% of PWS. We hypothesized that OA would have higher GCOW and net costs of walking (NCOW) at all speeds representative of walking speeds typical in daily life.

## 2. Methods

### 2.1 Participants

Twenty-eight participants were enrolled; one participant was excluded from analysis due to excessive talking during trials which distorted the metabolic measurements, leaving 27 participants for analysis (11 OA (6 Females (F)), 16 YA (7 Females(F))). This imbalance of number of participants between groups arose because younger adults were recruited both for the present comparison and as part of a broader study with different aims; rather than discard this additional data, we opted to include all available younger adult participants in our analysis. Sample size was not determined via a priori power analysis targeting a pre-specified effect size. Instead, our target sample size was informed by comparable prior studies examining the relationship between walking speed and metabolic cost, which have ranged from approximately 10 to 73 participants per age group (Larish et al., (1988): 17 older, 11 younger; Malatesta et al., (2003): 10 per group; Martin et al., (1992): 14–15 per age-by-activity group; Pearce et al., (1983): 22 older, 20 younger; Waters et al., (1988): 73 per group).

Exclusion criteria included chronic heart disease, diabetes, prior lower-limb surgery or joint replacements, neuromuscular injuries or falls (<6 months), or participation in specialized strength or endurance training. All participants provided written, informed consent, and the study was approved by the Delft University of Technology Human Research Ethics Committee. Participants were required to walk independently and perform daily activities without assistance. All participants followed standardized pre-test guidelines regarding fasting, alcohol, nicotine, and caffeine intake (Compher et al., 2006).

### 2.2 Protocol

After following manufacturer calibration guidelines, each participant was equipped with a COSMED K5 portable metabolic system (COSMED, Rome, Italy), a breath-by-breath metabolic analyzer. The system’s face mask covered the mouth and nose to capture breath-by-breath oxygen consumption*V̇O*_2_and carbon dioxide production*V̇CO*_2_. We also extracted walking cadence from the system using an embedded inertial measurement unit. We confirmed the protocol with the manufacturer to ensure appropriate calibration procedures and accuracy of measurements outdoors.

#### 2.2.1 Resting Trial

Prior to the outdoor walking trials, resting metabolic rate was measured while the participant stood unsupported and silently for seven minutes, also outdoors. Participants were instructed to stand still and visually focus on a single distant point to minimize head movement. Trials were conducted from summer to winter, with ambient air temperature ranging from 3 °C to 30 °C. Participants wore their own clothing, chosen for comfort at the prevailing ambient temperature, and skin temperature was not measured.

#### 2.2.2 Walking Trials

Participants completed eight, six-minute overground walking trials on an outdoor flat pavement course. The course consisted of a straight sidewalk, approximately 700m long, where participants walked back and forth for the duration of the trial, making U-turns at each end.

The first trial was conducted at PWS. Participants were instructed to walk at their, “usual, comfortable pace, without considering the goal of the destination.” PWS was determined from the distance walked in the six-minute trial, which was measured with a self-built pacing cart. The pacing cart comprised a Zozen collapsible measuring wheel to measure travelled distance and an N317 Retoo bike computer, enabling real-time speed monitoring.

The remaining trials were completed at fixed speeds in a randomized order: 0.8 m/s, 1.2 m/s, 1.6 m/s, PWS +5%, PWS -5%, PWS +10%, and PWS -10%. To regulate walking speed during the trials, participants were instructed to match the pace set by the pacing cart, carried by the researcher. To avoid influencing the step frequency and length, the researcher walked behind the participants, and a PVC pipe mounted on the wheel provided participants with a visual reference to monitor their speed independently. Between trials, participants were given up to five minutes to rest and hydrate, depending on their preference, and were invited to sit during these rest periods.

### 2.3 Data Processing and Analysis

Each sample of *V̇O*_2_ and *V̇CO*_2_ from the COSMED K5 system corresponded to a completed respiratory cycle, yielding an irregularly sampled time series with intervals at breathing frequency. For uniform temporal analysis, the breath-by-breath signals were linearly interpolated and resampled at 0.5Hz.

The cost of walking was defined as the metabolic energy expended per kilogram of body mass per meter traveled. Oxygen consumption was converted to metabolic cost with the following formulation (Brockway, 1987; WEIR, 1949).

GCOW and NCOW were calculated as follows (Equations 1 -3)

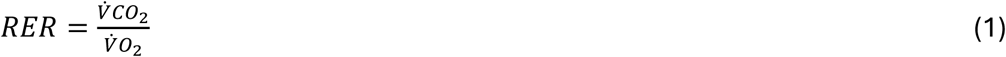

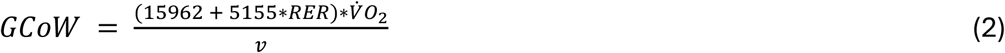

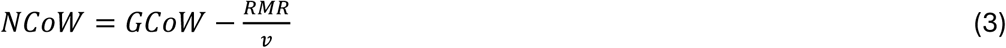

where RER is the respiratory exchange ratio, *V̇O*_2_ is the mass-normalized rate of oxygen consumption,*V̇CO*_2_ is the mass-normalized rate of carbon dioxide production, *v* is the average walking speed over the 6-minute trial, and RMR is the resting metabolic rate. Resting metabolic rate which was computed as the mean of mass-normalized metabolic power over the final 3 minutes of the 7-minute resting trial.

For each walking trial, we isolated a 3-minute steady-state window (See Supplementary Material for a representative plot of the time series data, Figure S7). Steady state was determined using a slope-based criterion applied to the resampled *V̇O*_2_ signal. The window with the smallest absolute slope was selected for analysis. We verified the slope of *V̇O*_2_; the maximum observed slope was 0.0027 L/min (M = 0.0001 L/min, SD = 0.0003 L/min). The mean GCOW and NCOW values computed within the selected window were used for analyses.

### 2.4 Statistical Analysis

A linear mixed-effects model was used to examine the effects of walking speed condition, age group, and their interaction on measured walking speed, cadence, NCOW, GCOW, and RER. Participant was included as a random intercept to account for repeated measures within individuals. Fixed effects included walking speed condition, age group, and their interaction. When a significant main or interaction effect was detected, post hoc pairwise comparisons were performed with Bonferroni correction to control for multiple comparisons (total *α*=0.05); for each pairwise comparison, the mean difference and corresponding 95% confidence interval (CI) were calculated alongside the Bonferroni-adjusted p-value. Differences in RMR between the young and older groups were assessed using an independent (unpaired) t-test (*α*=0.05).

## 3. Results

### 3.1 Participants

Younger adults were aged 23–30 years (n=16; mean ± SD: 26 ± 2 years; height: 172 ± 12 cm; body mass: 74 ± 12 kg). Older adults were aged 69–79 years (n=11; 74 ± 3 years; height: 173 ± 11 cm; body mass: 71 ± 10 kg). There were no significant differences in height or body mass between younger adults and older adults. Five individual trials were removed from analysis: one younger participant had three trials (0.8 m/s, 1.2 m/s, PWS - 10%) missing because testing was interrupted by rain, one younger participant had 1.2 m/s missing because the participant walked at a much lower speed initially, requiring a high acceleration during the trial, and one older participant had one trial (1.6 m/s) missing because they were unable to complete the task at the fastest walking speed. In the PWS trials across the cohort, we observed that participants regularly changed their speed throughout the trial. Therefore, MCOW outcomes from the PWS trials were excluded, and only the average speed from the PWS trial was retained for analysis.

### 3.2 Resting Metabolic Rate (RMR)

The RMR was 4.32 ml kg⁻¹ min⁻¹ for younger adults and 3.68 ml kg⁻¹ min⁻¹ for older adults. The RMR was not significantly different between groups, tested with an unpaired t-test (p=0.124). The older adult group contained one outlier data point, which contributed to the lack of statistical difference. We had no reason to exclude this outlier from our analysis, but their exclusion would result in a significant lower RMR in older adults compared to younger adults (p<0.01).

### 3.3 Walking speed and cadence

There was no significant main effect of age group in speed, (*F*(1, 195) = 0.08, *p* = .775) or cadence (F(1,195) =1.18, p=0.28), indicating that both groups walked at comparable speeds and cadences for all walking speed conditions, including at PWS (Table 1). For cadence, the age group effect at each individual condition was small and inconsistent in direction (range: −3.03 to +1.25 steps/min), with confidence intervals spanning roughly 9.5–13.6 steps/min. These intervals were large relative to the small observed group differences (1–3 steps/min at most conditions, Table 1, Figure 2). As designed, the conditions differed significantly from one another in speed (*F*(7, 196) = 223.68, *p* < .001), except for the PWS–10% (mean ± SD = 1.22 ± 0.11 m/s) and 1.2 m/s (1.21± 0.03 m/s) conditions, which did not differ significantly in speed (*p* = .146).

**Figure 1:**
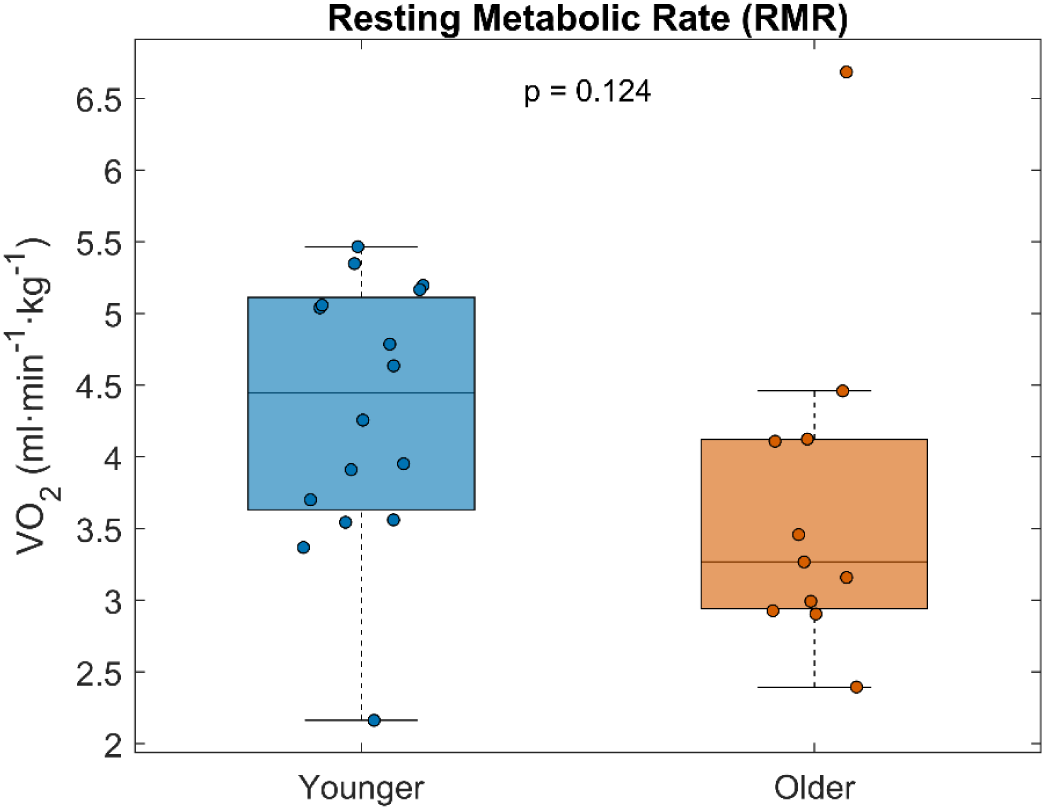
Resting metabolic rate (RMR) for younger (N=1C) and older (N=11) adult participants. There was not a significant difference between groups. The older adult group contained one outlier data point, which contributed to the lack of statistical difference. We had no reason to exclude this outlier from our analysis, but their exclusion would result in a significant lower RMR in older adults compared to younger adults (p**<**0.01).

**Figure 2:**
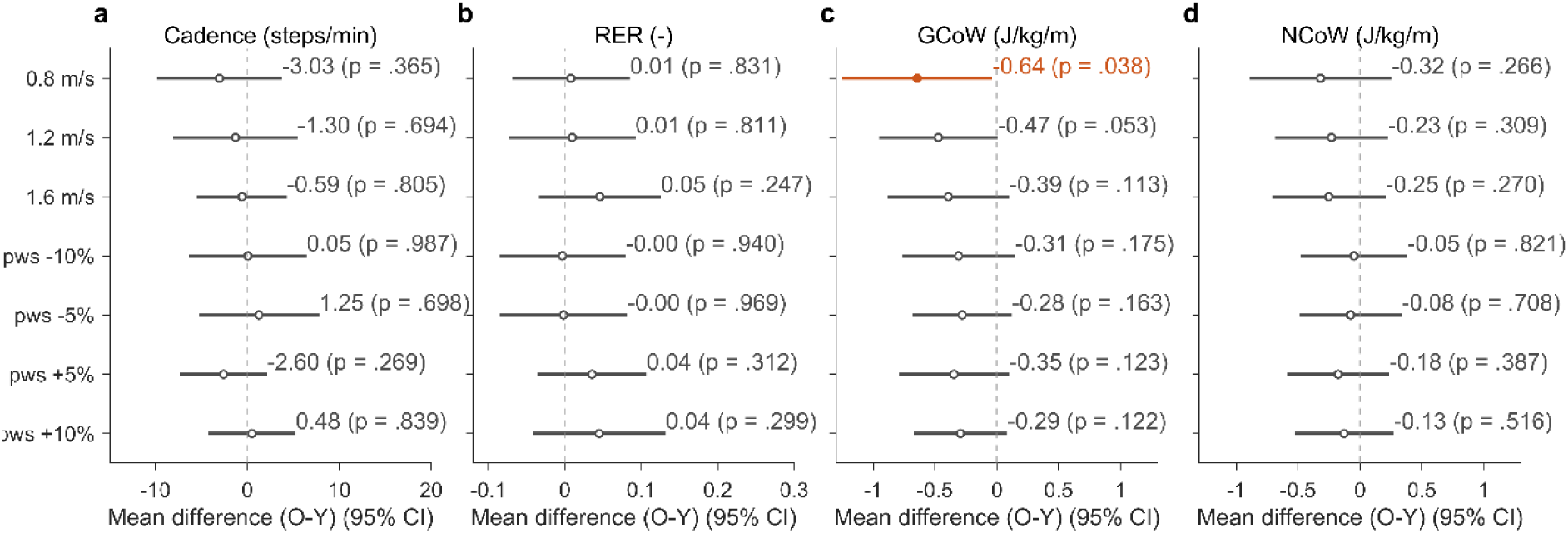
Pairwise comparisons between age groups (O:N=1C, Y: N=11) for each speed condition, expressed as mean differences with S5% confidence intervals (CI). Panels show (**a**) Cadence (steps/min), (**b**) RER (–), (**c**) GCoW (J·kg⁻¹·m⁻¹), and (**d**) NCoW (J·kg⁻¹·m⁻¹). Each row represents one pairwise comparison between age groups for seven speed conditions (0.8, 1.2, and 1.C m/s; preferred walking speed [PWS] −10%, −5%, +5%, and +10%). Dots indicate the mean difference between groups; horizontal lines indicate the S5% CI. The dashed vertical line at zero denotes no difference between groups. Comparisons for which the S5% CI does not cross zero (p < .05) are shown in orange; non-significant comparisons are shown in grey.

**Table 1.** Gait and metabolic parameters across walking speeds. * indicates a significant difference at 0.8 m/s between age groups (p<0.05).

| Parameter | Group | 0.8 | 1.2 | 1.6 | pws | pws_minus_10 | pws_minus_5 | pws_plus_5 | pws_plus_10 |
| --- | --- | --- | --- | --- | --- | --- | --- | --- | --- |
| n | Y | 15 | 14 | 16 | 16 | 15 | 16 | 16 | 16 |
|  | O | 11 | 11 | 10 | 11 | 11 | 11 | 11 | 11 |
| Speed (m s<sup>-1</sup>) | Y | 0.84 ± 0.04 | 1.21 ± 0.03 | 1.59 ± 0.04 | 1.35 ± 0.13 | 1.22 ± 0.11 | 1.28 ± 0.11 | 1.40 ± 0.13 | 1.49 ± 0.14 |
| | O | 0.85 ± 0.05 | 1.23 ± 0.04 | 1.58 ± 0.02 | 1.41 ± 0.13 | 1.28 ± 0.12 | 1.34 ± 0.12 | 1.46 ± 0.15 | 1.54 ± 0.14 |
| Cadence (steps min<sup>-1</sup>) | Y | 88 ± 11 | 113 ± 9 | 122 ± 6 | 117 ± 4 | 114 ± 7 | 115 ± 8 | 120 ± 5 | 120 ± 5 |
| | O | 85 ± 3 | 112 ± 8 | 121 ± 6 | 116 ± 7 | 114 ± 9 | 116 ± 9 | 118 ± 8 | 121 ± 8 |
| RER (–) | Y | 0.85 ± 0.10 | 0.84 ± 0.09 | 0.89 ± 0.09 | | 0.87 ± 0.11 | 0.86 ± 0.11 | 0.85 ± 0.09 | 0.86 ± 0.11 |
| | O | 0.86 ± 0.10 | 0.85 ± 0.11 | 0.93 ± 0.12 | | 0.87 ± 0.10 | 0.86 ± 0.11 | 0.89 ± 0.09 | 0.91 ± 0.12 |
| GCoW (J kg<sup>-1</sup> m<sup>-1</sup>) | Y | 3.77 ± 0.64* | 3.28 ± 0.62 | 3.35 ± 0.49 | | 3.21 ± 0.57 | 3.24 ± 0.58 | 3.25 ± 0.57 | 3.30 ± 0.54 |
| | O | 3.13 ± 0.92* | 2.81 ± 0.57 | 2.96 ± 0.78 | | 2.91 ± 0.59 | 2.96 ± 0.40 | 2.90 ± 0.58 | 3.00 ± 0.39 |
| NCoW (J kg<sup>-1</sup> m<sup>-1</sup>) | Y | 1.96 ± 0.77 | 2.02 ± 0.67 | 2.42 ± 0.57 | | 1.96 ± 0.58 | 2.09 ± 0.65 | 2.20 ± 0.59 | 2.30 ± 0.61 |
| | O | 1.64 ± 0.67 | 1.79 ± 0.40 | 2.17 ± 0.59 | | 1.91 ± 0.49 | 2.02 ± 0.29 | 2.02 ± 0.43 | 2.17 ± 0.34 |
| GCoW (ml kg<sup>-1</sup> m<sup>-1</sup>) | Y | 0.19 ± 0.03* | 0.16 ± 0.03 | 0.16 ± 0.03 | | 0.16 ± 0.03 | 0.16 ± 0.03 | 0.16 ± 0.03 | 0.16 ± 0.03 |
| | O | 0.15 ± 0.05* | 0.14 ± 0.03 | 0.14 ± 0.04 | | 0.14 ± 0.03 | 0.15 ± 0.02 | 0.14 ± 0.03 | 0.15 ± 0.02 |
| NCoW (ml kg<sup>-1</sup> m<sup>-1</sup>) | Y | 0.10 ± 0.04 | 0.10 ± 0.03 | 0.12 ± 0.03 | | 0.10 ± 0.03 | 0.10 ± 0.03 | 0.11 ± 0.03 | 0.11 ± 0.03 |
| | O | 0.08 ± 0.03 | 0.09 ± 0.02 | 0.10 ± 0.03 | | 0.09 ± 0.02 | 0.10 ± 0.01 | 0.10 ± 0.02 | 0.11 ± 0.02 |

PWS, preferred walking speed; RER, respiratory exchange ratio; GCoW, gross cost of walking; NCoW, net cost of walking; Y, young; O, older.

### 3.4 Respiratory Exchange Ratio (RER)

RER differed across walking speed conditions with a significant main effect (*F*(7, 195) = 7.29, *p* < .001). The main effect of age group was not significant (F(1,195) = 0.04, p = 0.84), indicating similar RER values for younger and older adults across walking speed conditions. Confidence intervals on the age group effect were narrow and centered near zero at every condition (±0.14–0.17; range: −0.003 to 0.046; Figure 2), indicating sufficient precision to interpret this as absence of a meaningful age effect. The largest confidence interval occurred at 1.6 m/s and PWS+10% (0.045–0.046), suggesting that an age-related difference, if present, may emerge at higher intensities. The walking speed condition × age group interaction was also not significant (F(7,195) = 1.31, p = 0.25), indicating that the effect of condition on RER did not differ by age group.

Bonferroni-corrected pairwise comparisons on the pooled participants showed that RER was significantly higher at 1.6 m/s compared to the other walking speeds (Figure 3, *p* < .05), except for PWS+10% which was not significantly different (*p* = .107). In addition, RER at PWS+5% was significantly higher than 1.2 m/s (*p* = .032) and significantly lower than 1.6 m/s (*p* = .004).

**Figure 3:**
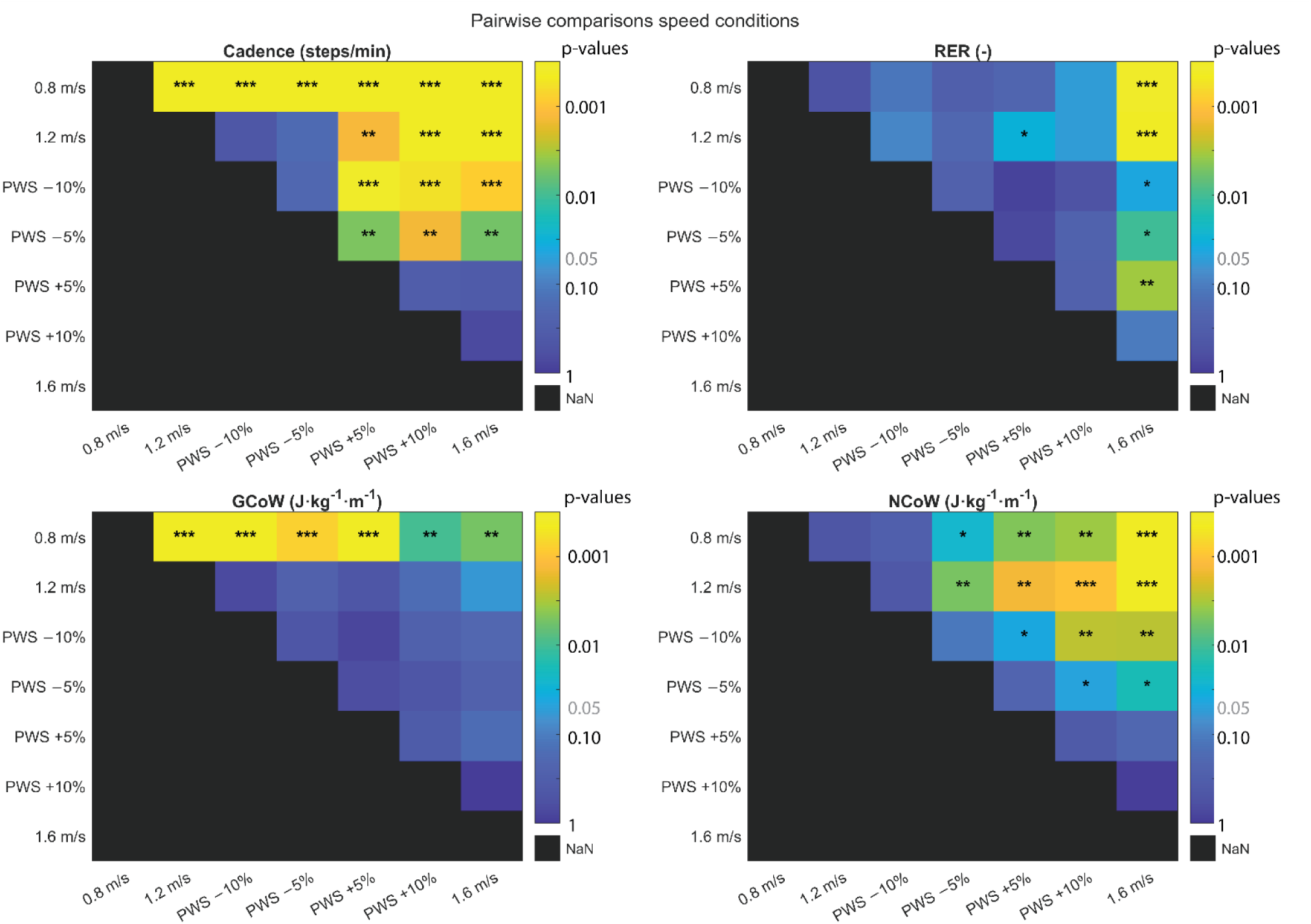
Statistical results from pairwise comparisons for cadence, respiratory exchange ratio (RER), gross cost of walking (GCOW), and net cost of walking (NCOW) across walking speed conditions. Pairwise comparisons (Bonferroni correction) are from the pooled sample of all participants (N = 27). Colors represent p-values on a logarithmic scale, from yellow (p < .001) to dark blue (p = 1); black cells indicate comparisons not applicable (NaN, diagonal only). Asterisks denote significance level: *p < .05, **p < .01, ***p < .001.

### 3.5 Gross Cost of Walking (GCOW (J kg⁻¹ m⁻¹))

There was a significant main effect of walking speed condition (F(7,195) = 7.35, p < 0.001), indicating that GCOW differed across walking speeds (Figure 4). There was also a significant main effect of age group (F(1,195) = 8.29, p = 0.004), indicating differences between younger and older adults in GCOW. The walking speed condition × age group interaction was not significant (F(7,195) = 1.34, p = 0.23).

**Figure 4:**
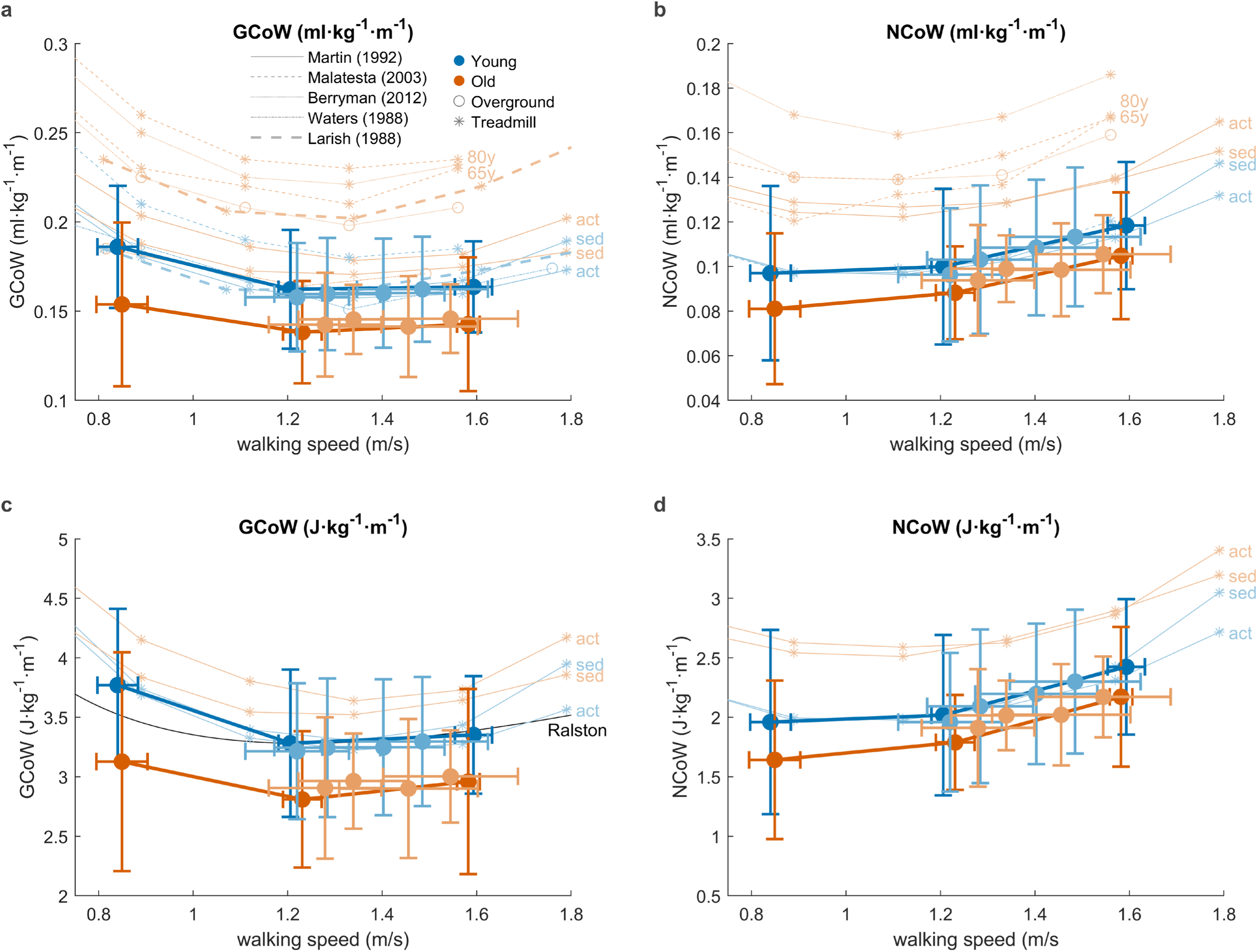
Gross cost of walking (GCOW) and net cost of walking (NCOW) across walking speed conditions for younger (N=1C) and older (N=11) adults. Data are presented in both ml•kg-1•m-1 (panels a and b) and J•kg-1•m-1 (Panels c and d). Present study data are plotted with that of Martin (P. E. Martin et al., 1SS2), Malatesta (2003), Berryman (2012), Waters (1S88), and Larish (1S88). Treadmill studies are indicated with * and overground studies are indicated with O. The hyperbolic fit from Ralston (1S58) is also plotted for GCOW (c). Data from Malatesta (2003) include older adult groups aged C5 and >80 yr. Data from Martin (1SS2) include sedentary (sed) and active (act) groups of younger and older adults.

Pairwise comparisons between age-groups revealed a significant difference at 0.8 m/s only, with older adults having lower GCOW than younger adults (OA: 3.13, SD = 0.92 J kg⁻¹ m⁻¹; YA: 3.77, SD = 0.64 J kg⁻¹ m⁻¹; p = 0.04; Figure 2); the groups were not significantly different at any other speed. However, the age group effect was negative for all seven conditions (range: −0.28 to −0.64 J kg⁻¹ m⁻¹, Figure 2). Confidence intervals at speeds other than 0.8 m/s did not exclude zero, but had similar magnitudes to the significant 0.8 m/s result (e.g., 1.2 m/s: 95% CI −0.95 to 0.01, p = 0.053; Figure 2). The consistent confidence intervals across independent conditions suggests our sample was only partially powered to resolve an effect that may be present throughout the full speed range.

Pairwise comparisons across the pooled sample of younger and older adults revealed that GCOW at 0.8 m/s was significantly higher than all other walking speeds (Table 1, Figure 3, p<0.01). Beyond 0.8 m/s, confidence intervals for the remaining pairwise comparisons were comparatively narrow (widths of 0.13–0.23 J kg⁻¹ m⁻¹), and consistently included zero, indicating no differences in GCOW in this speed range (Supplementary Material, 1.2–1.6 m/s and all PWS-adjusted conditions, Figure S5). One exception was 1.2 m/s versus 1.6 m/s, which had a narrow interval that did not quite exclude zero (95% CI: −0.296, 0.008; p = .062), suggesting a modest rise in cost at the fastest speed that did not reach significance.

### 3.6 Net Cost of Walking (NCOW (J kg⁻¹ m⁻¹))

NCOW differed across speeds, with a significant main effect of walking speed condition (Figure 4, F(7,195) = 5.15, p < 0.001), but no significant effect of age group (F(1,195) = 3.35, p = 0.069), and no walking speed condition × age group interaction (F(7,195) = 1.02, p = 0.42). The age group effect was negative at all seven conditions (Figure 2, range: −0.05 to −0.32 J kg⁻¹ m⁻¹), consistent in direction with the GCOW findings reported above, though confidence intervals were considerably wider than for GCOW and none reached significance individually (e.g., 0.8 m/s: 95% CI −0.89 to 0.26, p = 0.27). This wider uncertainty is expected given that net cost estimates combine the variance of both the loaded and resting metabolic measurements.

NCOW tended to increase with walking speed, but not between every walking speed condition (Figures 3 and 4). Bonferroni-corrected pairwise comparisons on the pooled sample showed that for NCOW, values at 0.8 m/s, 1.2 m/s, and PWS-10% did not differ significantly from each other, and all three were lower than the remaining conditions (Figure 3; p < 0.001–0.041). PWS-5% did not differ from its adjacent conditions, PWS- 10% (p = 0.113) or PWS+5% (p = 0.234), but was higher than the lower-speed conditions (0.8 m/s and 1.2 m/s; p < 0.025) and lower than the higher-speed conditions (PWS+10% and 1.6 m/s; p < 0.047). PWS+5%, which did not differ from PWS-5%, also did not differ from PWS+10% or 1.6 m/s. The two highest-speed conditions (PWS+10%, 1.6 m/s) had significantly higher NCOW than all other lower-speed conditions. Confidence intervals for NCOW pairwise comparisons were generally wider than for GCOW (widths of approximately 0.21–0.51 vs. 0.13–0.23 J kg⁻¹ m⁻¹), consistent with the greater variance inherent to NCOW estimates (Supplementary Material, Figure S5).

## Discussion

We hypothesized that older adults would have a higher MCOW than younger adults, both in GCOW and NCOW, at all walking speeds. This hypothesis was not supported at any of the seven walking speeds; rather, we observed a consistent trend toward slightly lower GCOW and NCOW in older adults, reaching significance for GCOW at 0.8 m/s only. While our sample size was modest, given the consistency of the direction of effect across all seven conditions and two related outcome measures (NCoW, GCoW), we would not expect a larger sample drawn from a similar population to reverse this trend. Our results indicate that, at matched overground walking speeds, our older cohort do not incur higher MCOW than younger adults.

A natural question is whether the older adults in our study were truly representative of the general older population. We did not administer a direct fitness assessment, so we relied on indirect comparisons. We expected to observe a slower PWS in our older adults (74 ± 3 yr), as previous studies have reported declines in PWS after 65 years of age (Schrack et al., 2012), which was not shown. PWS did not differ between age groups, which may suggest higher-than-average fitness in our older cohort. However, our older adults’ overground PWS (1.41 ± 0.13 m/s) aligns with the overground gait speed reported for older adults in a recent study (1.37 ± 0.15 m/s on flat terrain), which was notably higher than their treadmill-based estimate for the same participants (1.04 ± 0.20 m/s) (Santamaria-Guzman et al., 2026) . To further investigate, we compared our participants’ RMR and RER to prior literature distinguishing active and sedentary older adults. RMR trended lower in our older adults, and was significantly lower when excluding one outlier participant (Figure 1, p<0.01), consistent with prior work (Berryman et al., 2012; Frisard et al., 2007; Hernandes Júnior et al., 2021; P. E. Martin et al., 1992). Large-scale evidence shows that resting/basal energy expenditure declines from approximately age 60 onward (Pontzer et al., 2021), and physical activity/fitness status, independent of age, has been positively associated with RMR (Poehlman et al., 1991). Thus, the RMR data from our cohort do not support a higher-than-expected fitness level. The RER results show a similar outcome: in the younger adults, RER at 0.8, 1.2, and 1.6 m/s (0.85, 0.87, 0.89) fell between young active (0.84, 0.88, 0.88) and sedentary (0.84, 0.87, 0.89) groups from Martin et al. (1992) at comparable speeds. In the older adults, our RER values (0.86, 0.85, 0.93) aligned more closely with the sedentary older group (0.82, 0.89, 0.95) than the active older group (0.82, 0.87, 0.88), particularly at higher speeds (Figure 5). Taken together, neither RMR nor RER values suggest that our older participants were unusually fit. As is common in the field, our sample may be more active than the broad, general older adult population as they are healthy, mobile, and have volunteered to participate in research. But the RMR and RER evidence suggests this potential selection effect was modest rather than pronounced, as our older cohort appears more comparable to sedentary than to active reference groups in prior work.

**Figure 5:**
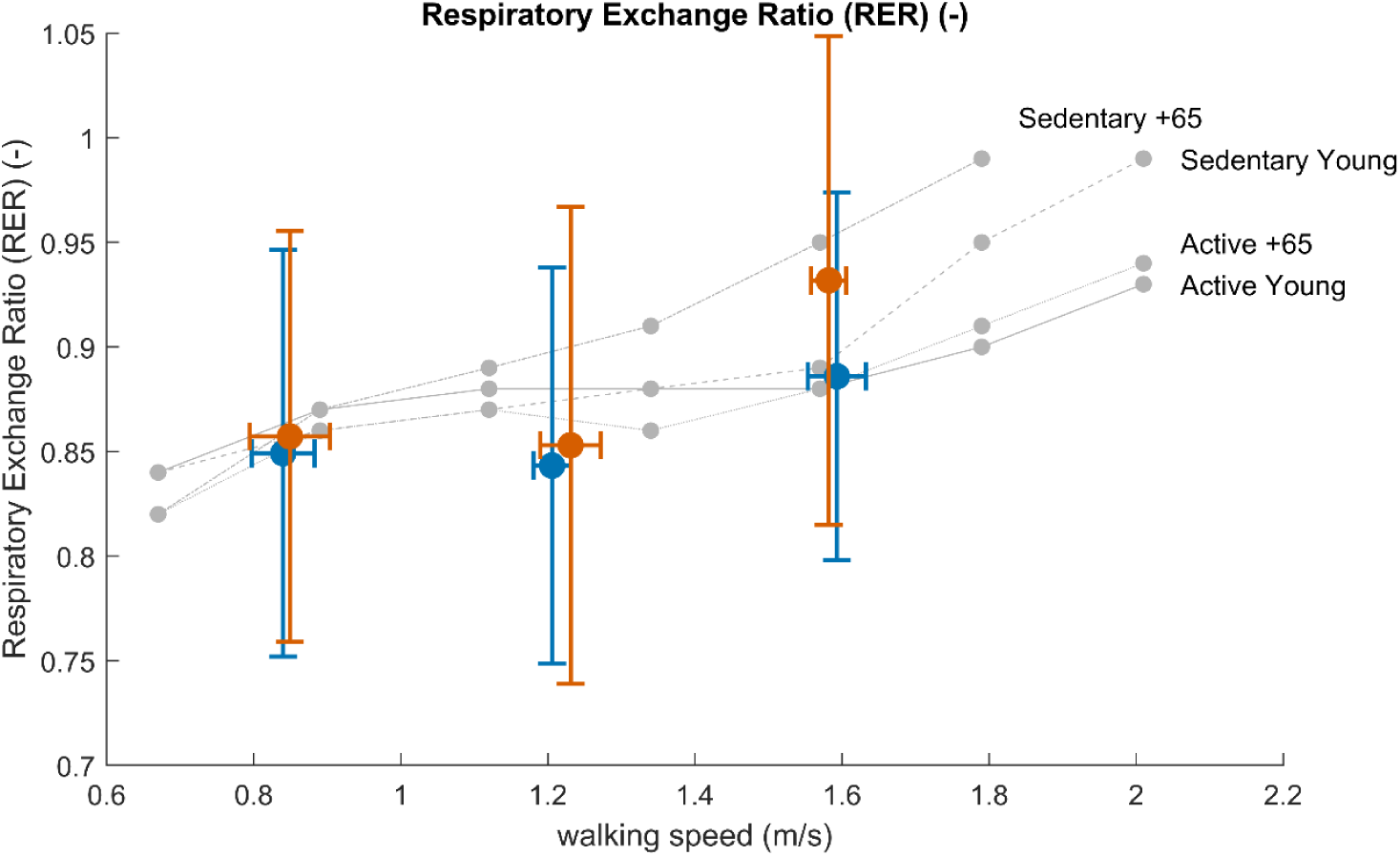
Respiratory exchange ratio (RER) across walking speeds for younger (blue, N=16) and older (orange, N=11) adults (mean ± SD), overlaid on reference data from Martin et al. (1SS2) for younger and older adults classified as active or sedentary (grey dashed lines). RER increased with walking speed in both age groups. Younger adults’ RER values fell between the active and sedentary young reference groups across all three speeds, while older adults’ values aligned more closely with the sedentary older reference group than the active older group, particularly at the higher speeds.

Our results contrast with treadmill-based studies reporting higher GCOW in older adults (Berryman et al., 2012; Malatesta et al., 2003; P. E. Martin et al., 1992; Zarrugh et al., 1974). This discrepancy may be partly explained by findings that older adults show a disproportionately greater increase in metabolic cost on a treadmill relative to overground walking compared with younger adults (Das Gupta et al., 2021). Notably, that same study found no age-group difference in overground cost at a single matched speed, consistent with a null result (Das Gupta et al., 2021). Our results extend this evidence by demonstrating that the absence of a higher MCOW in older adults persists across a range of overground speeds. Our results are also broadly consistent with earlier overground work by Pearce (1983) and Waters (1988), who also did not find higher MCOW overground in older adults, but their uncontrolled speed protocols precluded definitive confirmation, which is a limitation addressed by the present study. Differences between treadmill and overground findings in comparing older and younger adults, including our own, may partly reflect walking speed or context-dependent responses to the two settings rather than a fixed age-related difference in metabolic cost. However, we treat this as a tentative explanation, as our data cannot fully explain why the presence of an age effect would differ between settings. A direct, matched-speed comparison of overground and treadmill walking in younger and older adults is needed to test this potential explanation.

Independent of age-related effects, GCOW differed little across walking speeds in the pooled sample: only 0.8 m/s differed significantly from every other condition, while 1.2 m/s, 1.6 m/s, and the four PWS-adjusted conditions did not differ significantly from one another (Figure 3, Table 1), aside from a non-significant trend toward higher cost at 1.6 m/s relative to 1.2 m/s (p = .062). Our results are broadly consistent with prior literature on walking speed and metabolic cost. The relationship is typically characterized as “U-shaped”, which is a smooth, parabolic curve with a single, well-defined minimum, following Ralston et al. (1958). This interpretation has motivated simulation studies to search for the one speed that minimizes metabolic cost. We argue this description does not best represent the shape of the GCOW-speed curve shown in this study within this velocity range. Rather, GCOW is better described as a hyperbola with a flat range of speeds sharing statistically indistinguishable, near-minimal cost, rather than a smooth curve with one identifiable minimum. We agree that cost rises outside this flat region, which is precisely why the relationship is often, and understandably, characterized as U-shaped, but the presence of a near flat region is an important observation. The U-shape characterization traces largely to Ralston et al. (1958) whose U-shape curve was derived from a single individual with four data points; the pooled group data in their study (12M/7F, ages 22–51) presented a shallower, hyperbolic relationship, with cost, nearly flat between 1.1 and 1.4 m/s, closely matching our own results. (Figure 4). A hyperbolic group-level curve can, in principle, emerge from averaging individually parabolic curves with varying minima; while we could identify individuals in our dataset who show a similar U-shaped pattern in GCOW (Supplementary Material, Figure S2 for individual curves), most do not. Note that the two “arms” of the hyperbolic GCOW-speed curve are mostly driven by fundamentally different mechanisms: the left side, at low speeds (<1 m/s), is primarily shaped by RMR, while the right side, at higher speeds, reflects increases in mechanical work with velocity, as captured by NCOW. A truly parabolic, U-shaped relationship would require these two arms to be symmetric and governed by a shared underlying mechanism, yet they arise from two distinct physiological processes without producing a symmetric curve around a single minimum.

The importance of this flat region is strengthened by considering walking speeds outside the laboratory. The steep rise typically reported at very slow (<0.8 m/s) and fast (>1.6 m/s) speeds (Berryman et al., 2012; P. E. Martin et al., 1992; Ralston, 1958) falls outside the range commonly observed in natural, unconstrained walking among healthy adults. Our velocity ranges, PWS and cadence data align closely with Finley and Cody (1970), who observed 1,106 pedestrians outdoors in a natural environment (Figure 6). PWS in our cohort matched their male distribution (heights in our younger (172 ± 12 cm) and older (173 ± 11 cm) groups approximate U.S. male averages from 1971–1974 (175.26 cm vs. 161.54 cm for women (Abraham et al., 1979)), explaining the alignment with male walking speeds). Speeds below 0.8 m/s or above 1.6 m/s were practically absent in their large sample (Figure 6). Furthermore, Schrack et al. (2012) assessed PWS over 6-m indoors in 420 adults aged 32–96 years and speeds below 0.8 m/s were a small portion. Given that short indoor tracks likely underestimate outdoor, real-world walking speeds (Finley & Cody, 1970), speeds <0.8 m/s appear uncommon in adults. Velocities outside the 0.8–1.6 m/s range likely represent different modes of walking. Extremely slow speeds evoke a series of gait initiation events with different balance control demands. Extremely fast speeds approach or exceed the walk to run transition speed, suggesting that muscles are operating at the extremes of the force-length curve and not efficiently producing force (Neptune & Sasaki, 2005). These findings suggest that our study has effectively captured walking speeds that are common in community environments. In addition, our results show a higher GCOW at the low end of this range of speeds, which was similar for both age groups.

**Figure 6:**
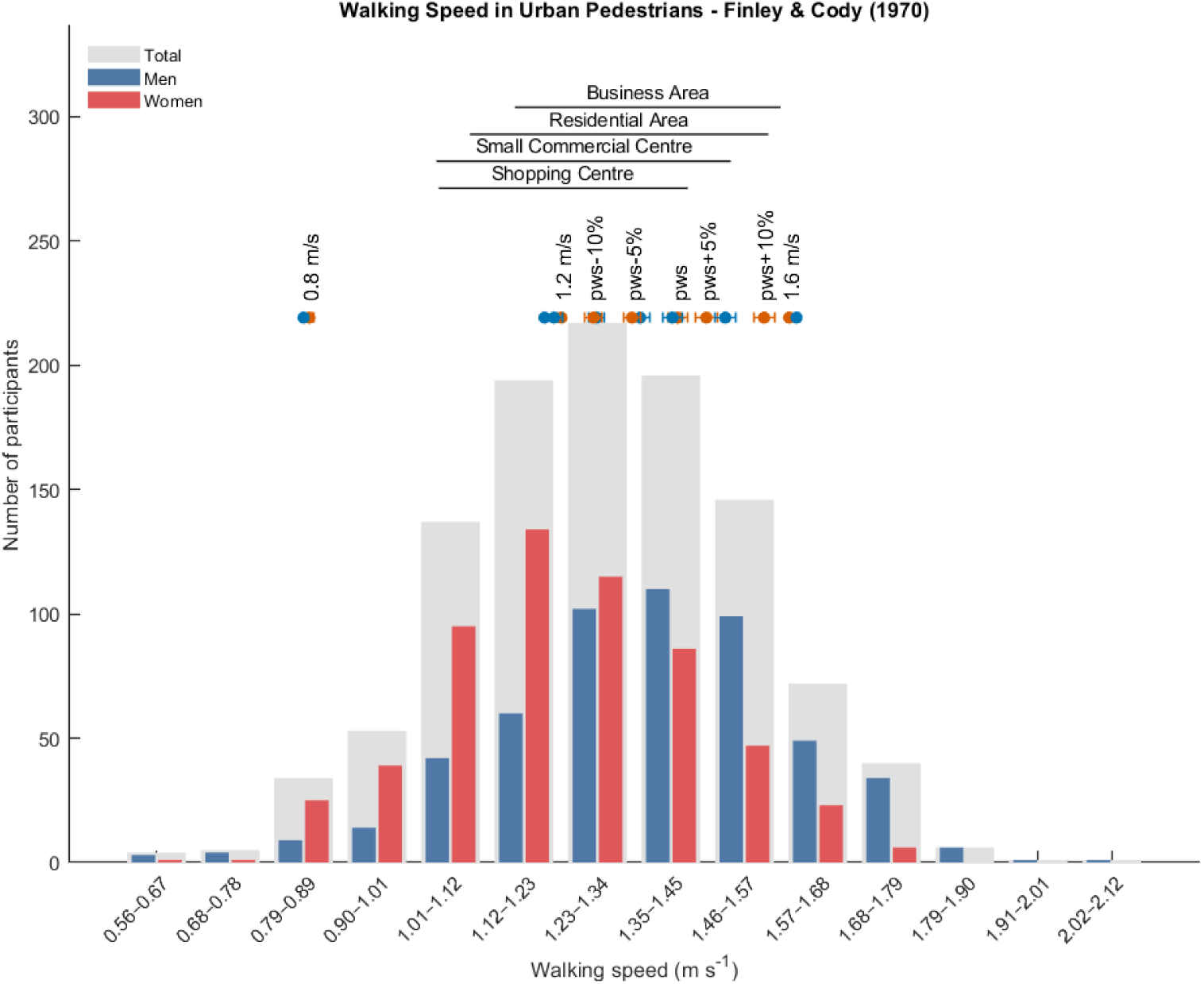
Histogram created from data from Finley and Cody (1S70). In their observational study, 1,10C pedestrians were unobtrusively observed in different areas of the city, and gait velocity and cadence were measured. Walking speed varied depending on the city district. The mean values with standard deviations from our study participants are displayed above the bars. Our PWS values are consistent with those reported for male pedestrians in their study. Walking speeds below 0.8 m/s and above 1.C m/s were rare.

NCOW increased with speed: the slowest conditions (0.8 m/s, 1.2 m/s, PWS−10%) did not differ from one another but were each significantly lower than the faster conditions (1.6 m/s, PWS+10%, PWS+5%), which also did not differ among themselves. This monotonic increase is consistent with prior literature reporting that net cost of walking rises with speed (Figure 3). This pattern was different from GCOW because NCOW isolates the metabolic work above resting rate (RMR). NCOW necessarily increases with mechanical demand, while GCOW includes the resting component that is proportionally larger at low speeds. In fact, the rise in GCOW at the low end of the speed range (0.8 m/s), is itself largely attributable to this resting component: because RMR is divided by speed to obtain its contribution to GCOW, this term is inflated at slow speeds (less distance covered per unit time) and shrinks rapidly as speed increases (>1m/s), even though RMR itself does not change with walking speed. This resting component also explains why an age group difference emerged for GCOW but not NCOW at 0.8 m/s. Because RMR is subtracted when computing NCOW, the component of GCOW that shows a difference between age groups is removed.

We acknowledge there are limitations that should inform future studies aiming to confirm or extend these findings:

- **PWS trials were not pace-controlled.** Because PWS was the only condition without pacing-cart regulation and participants frequently adjusted their speed within the trial. These accelerations inflated MCOW values in multiple participants, making them incomparable to constant-speed trials (Supplementary Material, Figures S2-S4). We suspect a similar mechanism may explain results of Pearce et al. (1983), who reported higher MCOW during overground (uncontrolled speed) versus treadmill (controlled speed) walking. Nonetheless, seven additional walking speeds enabled robust characterization of MCOW across the tested range. Future studies should determine PWS during familiarization and repeat the trial at this fixed speed.
- **Analytical methods may not have fully captured trial-level metabolic dynamics.** Approaches that account for within-trial speed variability, such as integrating metabolic cost over the recorded speed profile rather than assuming a constant pace, may help analyze less tightly controlled trials. Data collection should also continue until participants return to their resting metabolic rate following each trial, allowing the full metabolic response (including any excess post-exercise recovery) to be captured and accounted for in the analysis.
- **Ambient temperature was not tightly controlled.** Testing took place outdoors in summer through winter (all temperatures above 3°C; younger adults: 15 ± 10°C, older adults: 12 ± 5°C), unlike the climate-controlled conditions typical of treadmill studies. To mitigate this limitation, a gas calibration procedure ensured that ambient-air did not affect instrument calibration. We also performed a linear mixed model that included ambient temperature as a covariate, which showed consistent main effects and no age group × temperature interaction (Supplementary Material, Figure S6). This result suggests the observed age differences were not substantially driven by temperature. Participants wore their own clothing, which was selected for personal comfort under the prevailing conditions, and skin temperature was not directly measured. As such, ambient temperature may not fully reflect individual-level thermal exposure, and its potential influence on metabolic cost cannot be entirely ruled out. Future overground studies should consider standardizing clothing, directly measuring skin or core temperature, or restricting testing to a narrower ambient temperature range to more directly assess thermal effects.
- **Fitness level of older adults was not directly assessed.** We inferred relative fitness indirectly via RMR and RER comparisons to prior literature (see Discussion) rather than a direct measure, which cannot fully rule out our older cohort being more active than the general older population. We recommend that future studies also include a direct fitness assessment, such as measures of isometric strength, functional mobility measurements, and lean body mass, to characterize participants.
- **Other drivers of outdoor cost of walking were not examined.** Beyond age, factors such as spatiotemporal gait parameters — step length, step width, step frequency, stance/swing time, and their variability — are established correlates of metabolic rate on a treadmill (Selinger et al., 2015) but remain untested overground.

## Conclusion

Contrary to our hypothesis, older adults did not show a higher metabolic cost of walking than younger adults overground and outdoors across a wide range of walking speeds. Gross cost was significantly lower in older adults compared to younger adults at the slowest speed (0.8 m/s), with a similar but statistically not significant trend at the other speeds tested. These results contrast to the higher cost typically reported in older adults in treadmill studies, suggesting that laboratory findings may not translate directly to overground walking. Independent of age, GCOW was significantly higher at 0.8 m/s than at the remaining speeds (ranging 1.2–1.6 m/s) for both groups, confirming that walking at slower speeds increases GCOW, consistent with prior research.

## Supplementary Material

**Figure S1.**
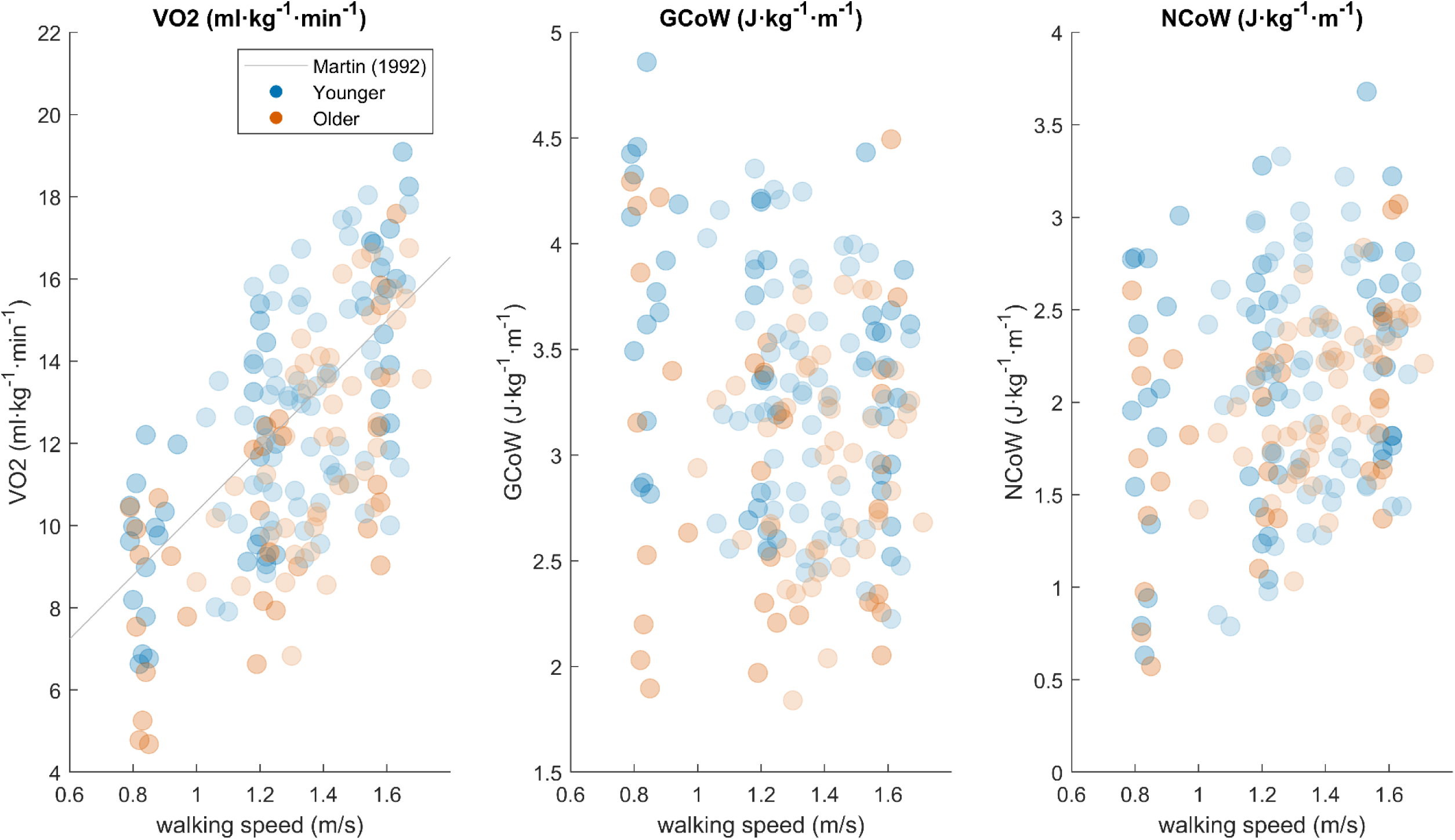
Individual data points for oxygen uptake (VO₂), gross cost of walking (GCOW), and net cost of walking (NCOW). The regression line reported by Martin et al. (1992) for measured VO₂ is overlaid for comparison. Data from younger (N=16) and older (N=11) participants are distinguished by color. YA = Younger Adults (Blue), OA = Older Adults (Orange)

**Figure S2.**
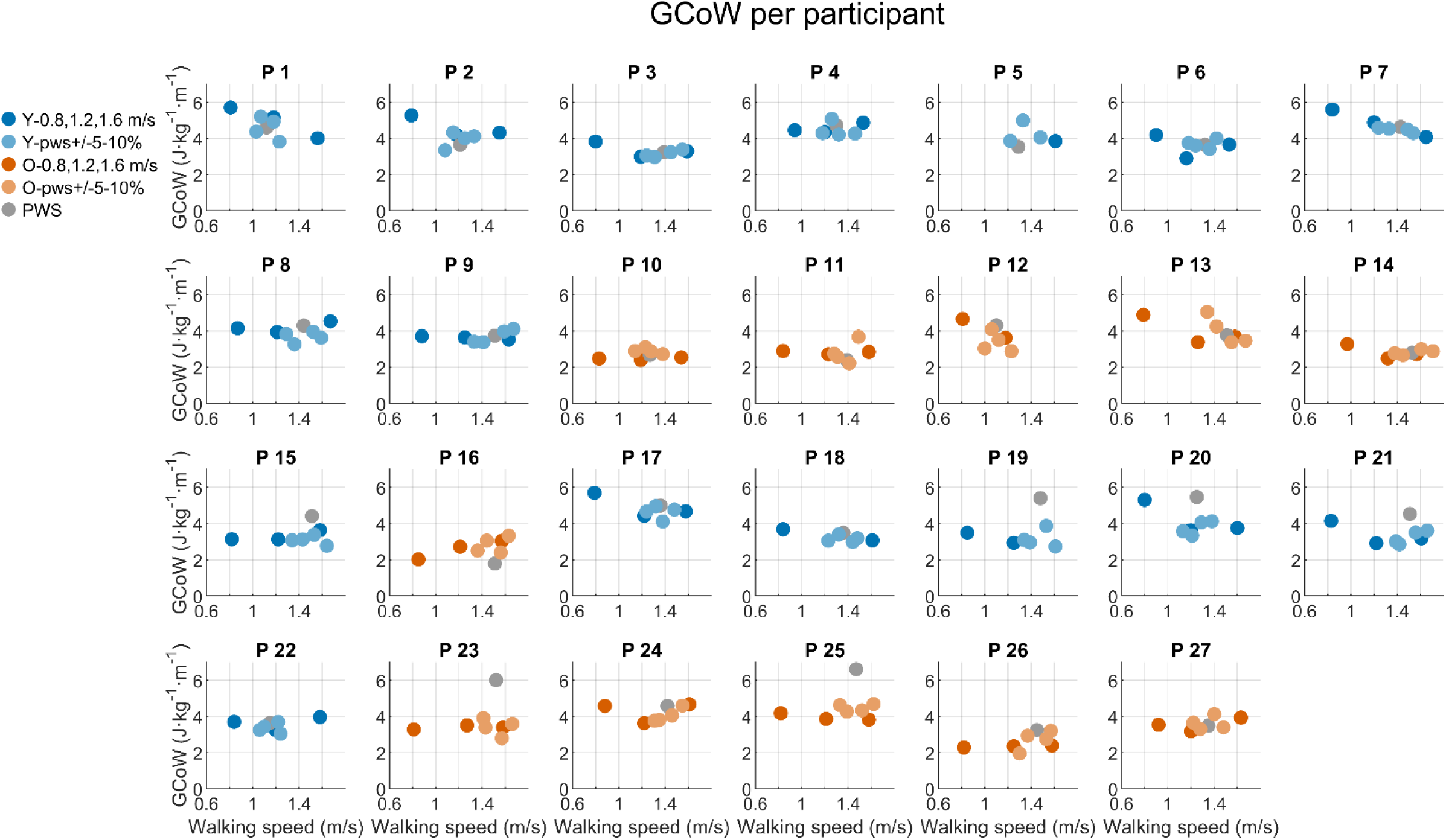
Gross cost of walking normalized to body weight and velocity as a function of walking speed for each individual participant.

**Figure S3.**
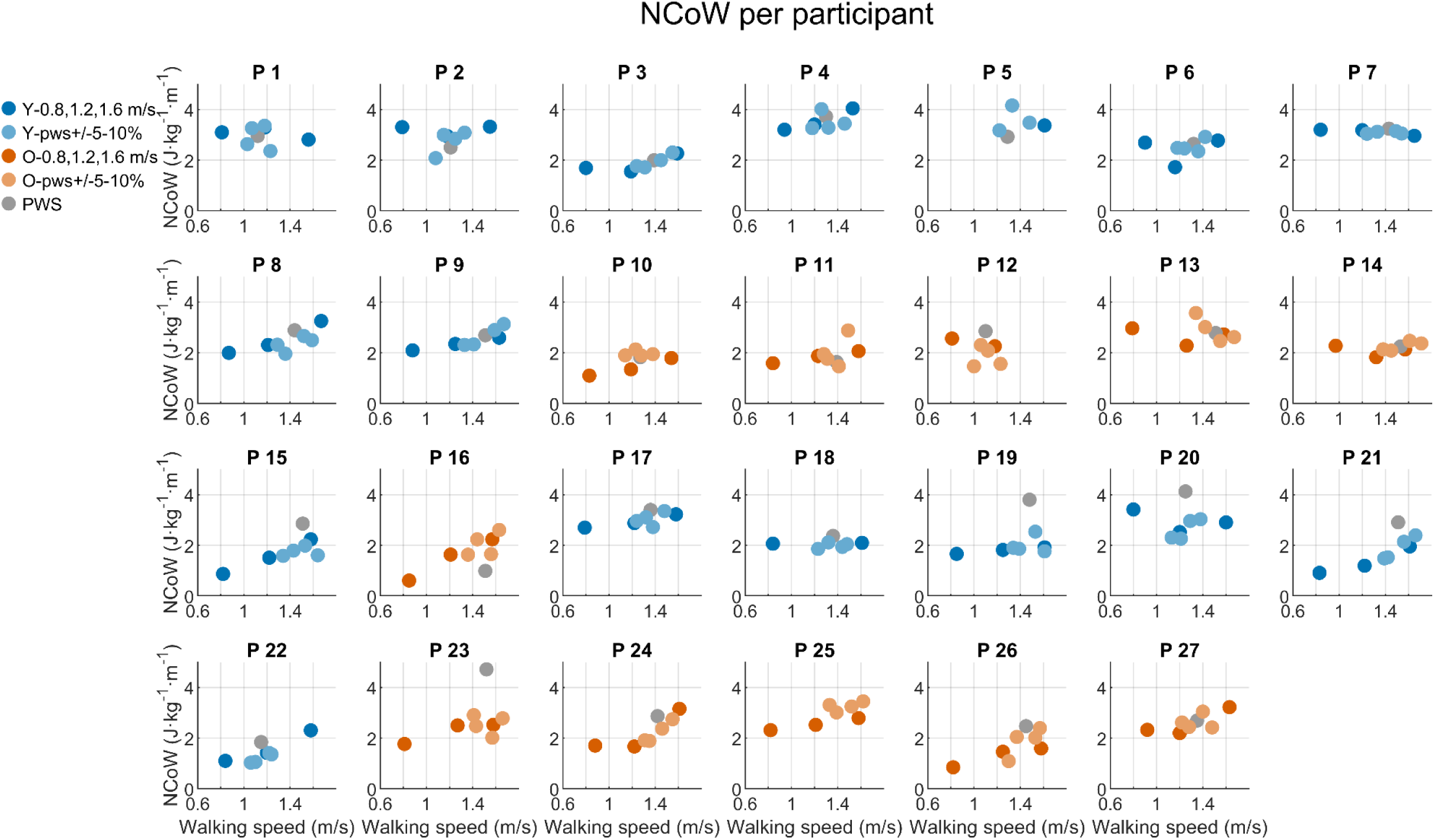
Net cost of walking normalized to body weight and velocity as a function of walking speed for each individual participant.

**Figure S4.**
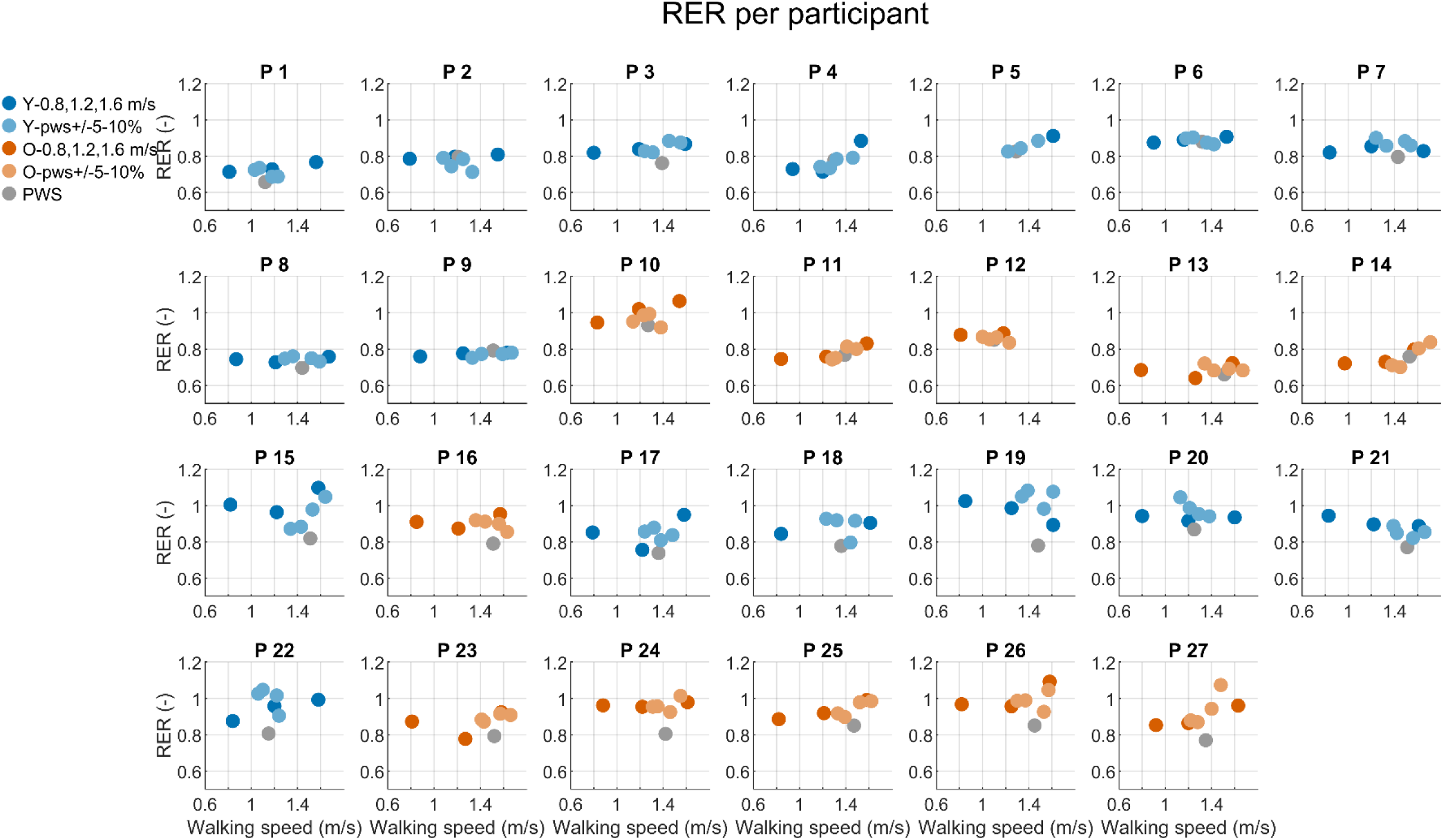
RER as a function of walking speed for each individual participant. Preferred walking speed (PWS) is indicated in grey. Solid

**Figure S5.**
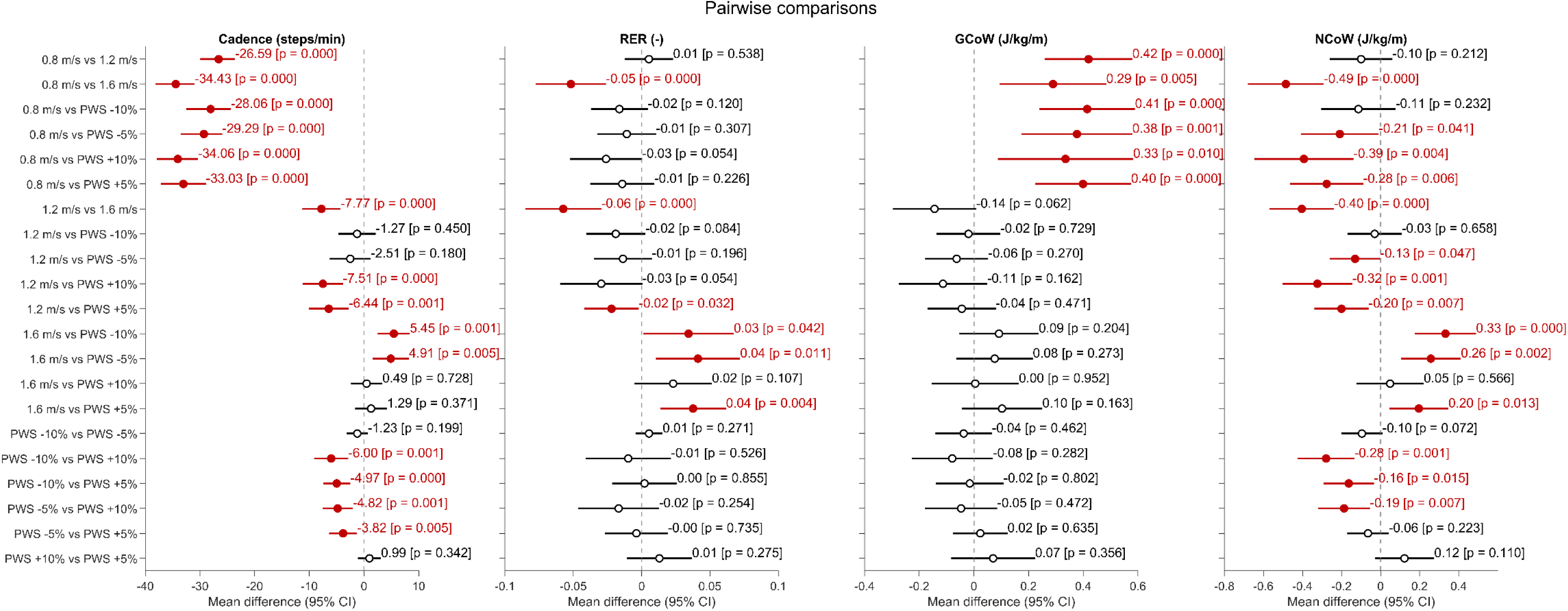
Pairwise comparisons between walking speed conditions across all participants (N=27) (**a**) Cadence (steps/min), (**b**) RER (–), (**c**) GCoW (J·kg⁻¹·m⁻¹), and (**d**) NCoW (J·kg⁻¹·m⁻¹), expressed as mean differences with 95% confidence intervals (CI). Each row represents one pairwise comparison between two walking speed conditions (0.8, 1.2, and 1.6 m/s; preferred walking speed [PWS] −10%, −5%, +5%, and +10%). Dots indicate the mean difference between conditions; horizontal lines indicate the 95% CI; the value and Bonferroni-adjusted p-value for each comparison are given alongside. The dashed vertical line at zero denotes no difference between conditions. Comparisons for which the 95% CI does not cross zero are shown in red; non-significant comparisons are shown in black.

**Figure S6.**
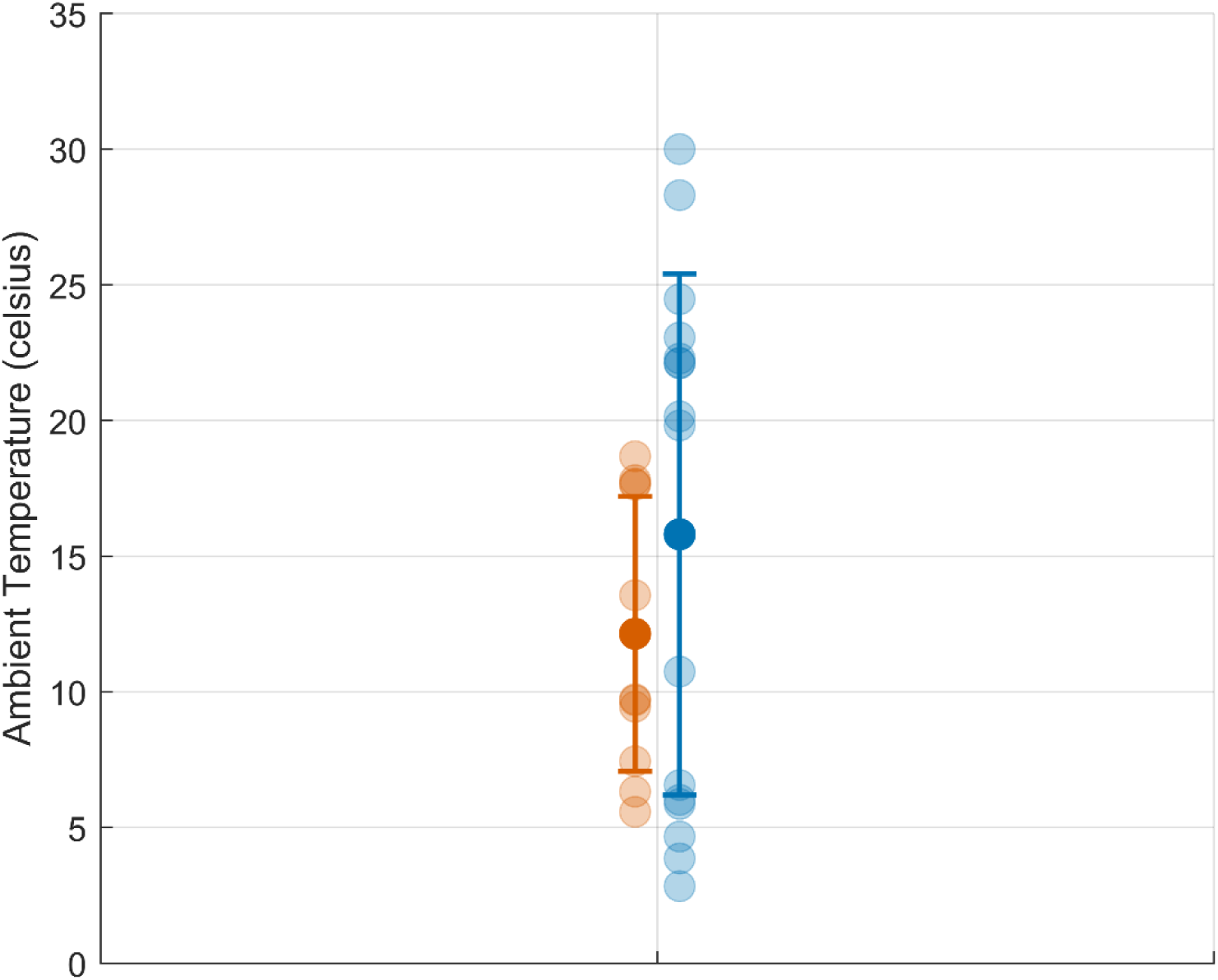
Ambient temperature (°C) recorded for Older adults (orange, N=11) and Younger adults (blue, N=16). Individual measurements are shown as semi-transparent dots, shifted horizontally for visibility. Filled circles indicate the mean for each group, with error bars representing ± standard deviation. Note that participants wore their own clothing, selected for their own comfort at the given ambient temperature; as actual skin temperature was not measured, ambient temperature may not fully reflect individual thermal exposure.

**Figure S7.**
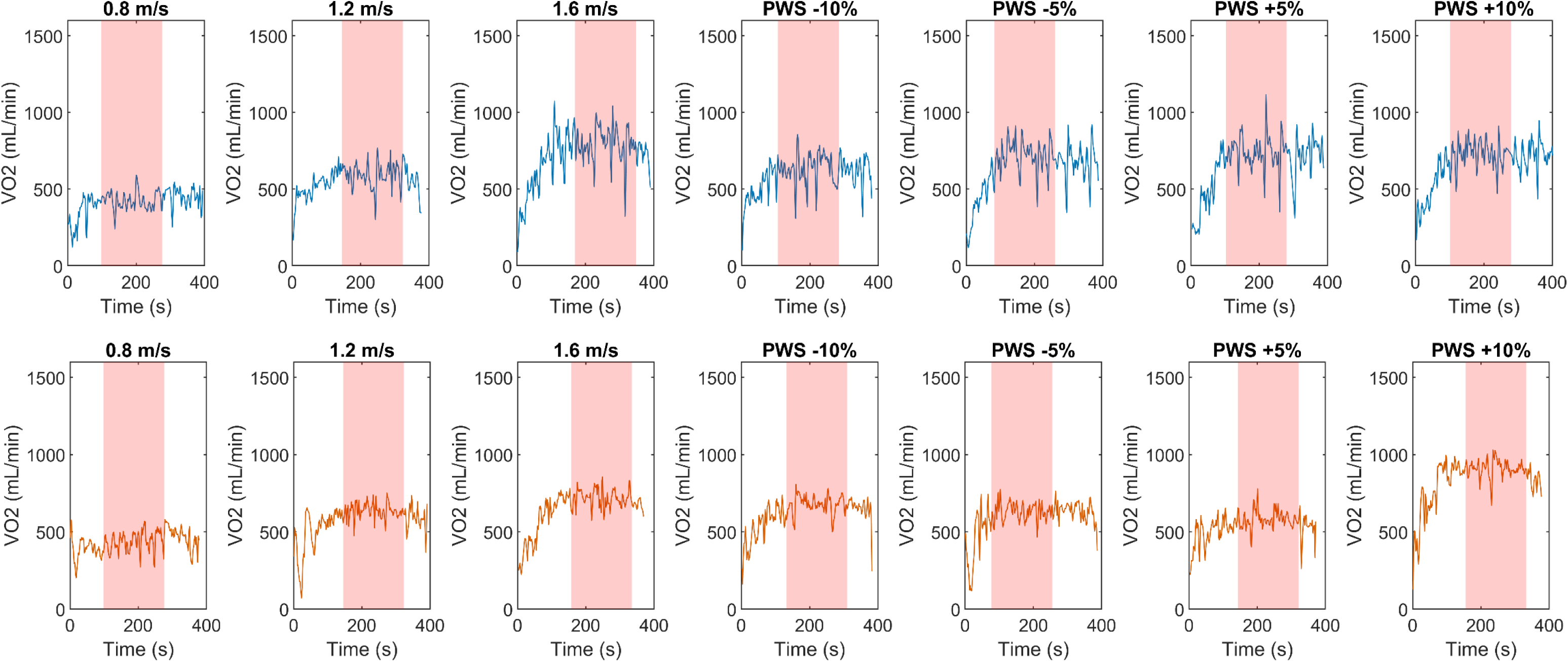
Breath-by-breath oxygen consumption (VO₂) time series across the seven walking conditions (0.8, 1.2, and 1.6 m/s fixed speeds, and preferred walking speed (PWS) perturbations of −10%, −5%, +5%, and +10%) for two representative participants: a younger adult (top row, P11, blue) and an older adult (bottom row, P15, orange). Shaded red regions indicate the steady-state window used for VO₂ estimation in each trial. VO2 rises from a low baseline at trial onset and plateaus within the shaded window, from which steady-state values were extracted for further analysis.

